# Brain microRNAs associated with late-life depressive symptoms are also associated with cognitive trajectory and dementia

**DOI:** 10.1101/620815

**Authors:** Thomas S. Wingo, Jingjing Yang, Wen Fan, Benjamin Logsdon, Se Min Canon, Bing Yao, Nicholas T. Seyfried, James J. Lah, Allan I Levey, Patricia A. Boyle, Julia A. Schneider, Philip L. De Jager, David A. Bennett, Aliza P. Wingo

## Abstract

**Objective:** Late-life depression is associated with an increased risk for dementia, but our knowledge of the molecular mechanisms underlying this association is limited. Hence, the authors investigated whether microRNAs, important post-transcriptional regulators of gene expression, contribute to this association.

**Method:** Late-life depressive symptoms were assessed annually in 300 non-demented participants of the Religious Orders Study and Rush Memory and Aging Project for a mean of seven years using the Center for Epidemiological Studies Depression scale. Participants underwent annual cognitive testing, clinical assessment of cognitive status, and uniform neuropathologic examination after death. microRNAs were profiled from the prefrontal cortex using Nanostring platform. A global microRNA association study of late-life depressive symptoms was performed using linear mixed model adjusting for sex, age, Alzheimer’s dementia pathological burden, proportions of brain cell types, post-mortem interval, and RNA integrity.

**Results:** Four brain microRNAs were associated with late-life depressive symptoms at adjusted p<0.05 (miR-484, miR-26b, miR-30d, and miR-197). Lower expressions of these miRNAs were associated with greater depressive symptoms. Furthermore, lower expressions of miR-484 and miR-197 were associated with faster decline of cognitive performance over time. Additionally, lower miR-484 level was associated with higher probability of having Alzheimer’s dementia. Lastly, the predicted targets of miR-484 were enriched in a brain protein co-expression module involving synaptic transmission and regulation of long-term neuronal synaptic plasticity.

**Conclusions:** This is the first study to identify brain microRNAs associated with late-life depressive symptoms assessed longitudinally. Additionally, the authors found a link between late-life depressive symptoms and dementia through miR-484 and miR-197.

---

Late-life depression is associated with an increased risk for dementia with an odds ratio of 1.85 according to meta-analyses of 23 prospective studies including more than 49,600 participants (1, 2). Notably, depression has been shown to be associated with faster decline of cognitive performance and with the clinical diagnosis of dementia but not with the traditional dementia pathologies such as beta-amyloid plaques, tau tangles, Lewy bodies, hippocampal sclerosis, microinfarcts, and macroinfarcts (3–6). How depression and its associated molecular changes act to increase dementia risk is not well understood and is the focus of this work.

MicroRNAs (miRNAs) are important post-transcriptional regulators of gene expression and have been implicated in many neuropsychiatric disorders (7–10). miRNAs are sometimes referred to as “master regulators” because one miRNA can regulate hundreds of genes and thus can exert a substantial effect on the gene expression networks (10). So far, two studies have examined brain miRNA profile in major depression with sample sizes of less than 35 (8, 11). Hence, studies of global miRNA profile in depression with a larger sample size and with a focus on late-life depression are needed to provide more generalizable findings and insights into mechanisms by which depression increases risk for dementia.

To understand the molecular mechanisms behind detrimental effect of depression on dementia risk, we first performed a global miRNA association study of longitudinally assessed depressive symptoms in non-demented older adults who were followed for a mean of seven years. We found four miRNAs significantly associated with late-life depressive symptoms. Next, we examined these four miRNAs in relation to the dementia-related traits, including rate of decline of cognitive performance over time, clinical diagnosis of mild cognitive impairment or dementia, and eight neuropathologies. Two miRNAs were found to be associated with these dementia features (miR-484 and miR-197). Then, we examined the targets of these two miRNAs in a published protein co-expression network and found that targets of miR-484 were enriched in the protein co-expression module involved in synaptic transmission and regulation of long-term synaptic plasticity. Together, these findings point to a molecular link between depression and elevated risk for dementia for further mechanistic studies.

## Methods

### Participants

Participants were from two longitudinal clinical-pathologic cohort studies of aging and Alzheimer’s disease - Religious Orders Study (ROS) and Rush Memory and Aging Project (MAP) (12). ROS is comprised of older Catholic priests, nuns, and monks throughout the USA. MAP recruits older lay persons from the greater Chicago area. Both studies involve detailed annual cognitive and clinical evaluations and brain autopsy. Participants provided informed consent, signed an Anatomic Gift Act, and a repository consent to allow their data and biospecimens to be repurposed. The studies were approved by the Institutional Review Board of Rush University Medical Center. To be included in our global miRNA association study of depressive symptoms, participants must not carry a final clinical diagnosis of dementia.

### Neuropsychiatric phenotypes

#### Depressive symptoms

were assessed annually using the 10-item version of the Center for Epidemiological Studies Depression scale (CES-D) (13). The CES-D captures the major symptoms of depression as identified in clinical and factor analytical studies, i.e. depressed affect, positive affect, somatic complaints, and interpersonal problems, with a reliability of 0.8 (13). The possible score range for the CES-D is 0 to 10, with higher score indicating more depressive symptoms.

#### Final clinical diagnosis of dementia

is determined at the time of death by a neurologist with expertise in dementia using all available clinical data but blinded to postmortem data. The diagnosis was based on the recommendation of the National Institute of Neurological and Communicative Disorders and Stroke and the AD and Related Disorders Association (14). Case conferences including one or more neurologists and a neuropsychologist were used for consensus as necessary. Clinical diagnoses of cognitive status can include no cognitive impairment, mild cognitive impairment (MCI), or Alzheimer’s disease (AD).

#### Rate of decline of cognitive performance

is the individual rate of decline of global cognitive performance over time. Annually, 21 cognitive tests were administered to each ROS/MAP participant with 17 in common. The raw scores from 17 cognitive tests were standardized to a Z score with respect to the mean and standard deviation of the cohort at the baseline visit. These Z scores were averaged to create the composite annual global cognitive score. Rate of cognitive change is the random slope with respect to follow-up years in the linear mixed model in which the annual global cognitive performance was the longitudinal outcome, follow-up year was the independent variable with random effect per subject, and age at recruitment, sex, and years of education were the covariates.

#### Dementia pathologies

Brain autopsy was performed by examiners who were unaware of deceased participants’ clinical information and have been described in detail before (15, 16). Nine brain regions of interest (i.e. midfrontal, midtemporal, inferior parietal, anterior cingulate, entorhinal and hippocampal cortices, basal ganglia, thalamus, and midbrain) were dissected and stained for assessment of pathology. Global AD pathology (i.e. neuritic plaques, diffuse plaques, and neurofibrillary tangles) was visualized in five cortical regions using a modified Bielschowsky silver stain. Counts of silver-stained neuritic plaques, diffuse plaques, and neurofibrillary tangles were used to create a continuous measure of AD global pathology. The square root of this global pathology measure was used in our analyses to improve its normal distribution. Chronic gross infarcts were identified visually by examining slabs and pictures from both hemispheres and confirmed histologically, and was treated as a dichotomous variable (present vs. absent) in our analyses. Microinfarcts were those that were not visible to the naked eye but were identified under microscope using haematoxylin and eosin stain in a minimum of nine regions, including six cortical regions, two subcortical regions, and midbrain. Microinfarcts were treated as present or absent in our analyses. Lewy body pathology was assessed using antibodies to α-synuclein in six regions including substantia nigra, limbic and neocortices, and treated as present or absent in our analyses. Hippocampal sclerosis was identified as severe neuronal loss and gliosis in hippocampus or subiculum using haematoxylin and eosin stain and treated as present or absent in analyses (17). TDP-43 cytoplasmic inclusions were assessed in six regions using antibodies to phosphorylated TDP-43. Inclusions in each region were rated on a six-point scale and the mean of the regional scores was created (18). TDP-43 scores were dichotomized into absent (i.e. mean score of 0) or present (mean score >0) in our analyses. Cerebral atherosclerosis was assessed by visual inspection of vessels in the circle of Willis and rated as absent, mild, moderate, or severe, and treated as a semiquantitative variable in our analyses (19). Cerebral amyloid angiopathy (CAA) was assessed using amyloid-β immunostaining in four regions (i.e. midfrontal, inferior temporal, angular, and calcarine). In each region, meningeal and parenchymal vessels were assessed for amyloid deposition and scored from 0-4, where 0 indicates no deposition, 1 refers to scattered segmental but no circumferential deposition, 2 means circumferential deposition up to 10 vessels, 3 reflects circumferential deposition in > 10 vessels and up to 75% of the vessels, and 4 indicates circumferential deposition in over 75% of the vessels (20). The score for each region was the maximum of the meningeal and parenchymal scores, and a continuous summary score was created by averaging scores across the regions (20).

### miRNA quantification and quality control

Quantification and quality control of miRNA profiles were described in detailed before (21). Briefly, total RNA was extracted from the dorsolateral prefrontal cortex (dlPFC). Then, small RNA was isolated from total RNA and profiled using the Nanostring platform. RNA integrity number (RIN) ranged from 5 to 10. Samples were randomized with respect to clinical diagnosis of cognitive status to minimize batch effects. miRNAs from the Nanostring RCC files were re-annotated using miRBase v19. After correcting for the probe-specific backgrounds, a three-step filtering of miRNA expressions was performed. First, miRNA with missing expression levels in > 95% of samples were removed. Second, samples with > 95% of miRNAs with missing expression were removed. Third, all miRNA whose absolute values were less than 15 in at least 50% of the samples were removed to eliminate miRNA that had negligible expression in brain samples. After the miRNA and sample filtering, the dataset consisted of 292 miRNAs. Combat was used to remove batch effects and quantile normalization was performed (22).

### Estimate of brain cell type proportions

The total RNA extracted from the dlPFC in ROS/MAP subjects was used for miRNA profiling as well as for RNA sequencing as described in detail before (23). We estimated the proportions of neurons, astrocytes, oligodendrocytes, and microglia for the dlPFC brain tissue from which RNA was extracted for each ROS/MAP subject. We used Darmanis’ single-cell RNA-sequencing expression profiles from human tissue samples as reference and deconvoluted RNA sequencing profiles for ROS/MAP subjects (24). We then used the proportions of these four cell types as covariates in all our miRNA analyses to control for the heterogeneity of cell types in brain tissues.

### Global miRNA association study of depressive symptoms

Using linear mixed model, we examined the association between longitudinally assessed depressive symptoms and global miRNA profile, adjusting for sex, age at visit, global AD pathology burden, proportions of neurons, oligodendrocytes, astrocytes, and microglia, post-mortem interval (PMI), RIN, and study (ROS or MAP). Specifically, depression score was modeled as the outcome, miRNA as a predictor, with a random intercept and random slope for each subject with respect to the follow-up year, adjusting for the above-mentioned covariates. In this linear mixed model, we tested whether the miRNA abundance had a non-zero effect on the depression scores. By using a mixed effect on follow-up year (a covariate in this model), we accounted for the individual-specific change in depression score from year to year in the association test. To address multiple testing, we used Benjamini-Hochberg (BH) method to control the false discovery rate, and declared significantly associated miRNAs as those with an adjusted p<0.05 (25).

### Depressive symptoms-associated miRNAs versus dementia features

We examined associations between depressive symptoms-associated miRNAs from the global miRNA analysis above and the dementia-related features. These dementia features include i) rate of decline of cognitive performance; ii) clinical diagnosis of dementia; and iii) neurodegenerative pathologies including global AD pathology, Lewy bodies, TDP-43, cerebral atherosclerosis, gross infarcts, chronic microinfarcts, cerebral amyloid angiopathy, and hippocampal sclerosis. We included all ROS/MAP participants with miRNA profiles in this analysis, hence the sample size for these analyses is greater than 300. We used general linear regression model in which specific dementia feature was the outcome, miRNA was the predictor, and sex, age at death, education, PMI, RIN, proportions of neurons, oligodendrocytes, astrocytes, and microglia, and study as covariates.

### Target genes of miR-484 and miR-197 and protein co-expressed modules

A published study identified 16 modules of co-expression proteins in the Baltimore Longitudinal Study of Aging (BLSA) (26). Since a miRNA represses expression of its target genes by either destabilizing the transcripts or repressing the translation of transcripts to proteins, the effects of miRNAs ought to be most apparent at the protein level. Hence, we asked whether the targets of miR-484 or miR-197 would be enriched in any of these protein co-expression modules. To that end, predicted targets of miR-484 and miR-197 were identified from TargetScan database v7.2 (27). Next, we asked whether these targets were enriched in any of the 16 modules of co-expressed proteins mentioned above using gene set enrichment analysis and adjusting for multiple testing with Benjamini-Hochberg (BH) method.

## Results

### ROS/MAP participants

A total of 300 non-demented ROS/MAP subjects were included in the global miRNA analysis of depressive symptoms assessed longitudinally. Table 1 lists the characteristics of the subjects. Briefly, these participants had an average of 16 years of education, 61% were women, 56% had normal cognition and 44% had mild cognitive impairment (Table 1). These participants were followed for a mean of seven years and a median of 6 years. Mean age at enrollment was 65 and mean age at death was 87. The depression score for each individual averaged over all the follow-up years ranged from 0 to 7.

**Table 1:** Descriptive characteristics of the ROS/MAP participants (N=300)

| **Characteristic** | **N** | **Percent** |  |
| --- | --- | --- | --- |
| Sex (Female) | 183 | 61.0 |  |
| Diagnosis rendered at death |  |  |  |
| Normal cognition | 167 | 55.7 |  |
| Mild cognitive impairment | 133 | 44.3 |  |
| Study |  |  |  |
| ROS | 164 | 54.7 |  |
| MAP | 136 | 45.3 |  |
|  | **Mean [SD]** | **Median** | **Range** |
| Age at enrollment | 65.3 [7.2] | 80.4 | [65 – 98] |
| Age at death | 86.9 [6.6] | 86.9 | [67 – 104] |
| Education | 16.5 [3.5] | 16.0 | [5 – 26] |
| Depression score |  |  |  |
| (average over all follow-up years) <sup>a</sup> | 1.3 [1.3] | 0.9 | [0 – 7] |
| Rate of cognitive decline <sup>b</sup> | 0.04 [0.04] | 0.04 | [-0.10 – 0.14] |
| Number of follow-up years | 6.9 [3.9] | 6.0 | [1 – 17] |
| Post-mortem interval (PMI) | 7.3 [5.0] | 5.9 | [1 – 32.6] |
| RNA integrity number (RIN) | 7.2 [1.0] | 7.4 | [5 – 9.9] |

<sup>a</sup> CES-D has a possible score of 0 to 10 with the higher score, the more depressive symptoms
<sup>b</sup> The more negative the value, the faster the decline

### Global miRNA association study of late-life depressive symptoms

We found four miRNAs significantly associated with late-life depressive symptoms at an adjusted p-value < 0.05 after accounting for sex, age, global AD pathology, cell type, PMI, RIN, and study (Figure 1, Table 2A, Supplementary Table 1). All four miRNAs (miR-484, miR-26b, miR-30d, and miR-197) were less abundant in persons with greater depressive symptoms (Table 2A). To understand whether these results depended on AD pathology found at the end of life, a separate model that did not adjust for AD pathology was used, which showed similar findings except that miR-30d had adjusted p of 0.06 instead of adjusted p value of 0.048. Together, these analyses suggest that these miRNAs are associated with depressive symptoms independent of AD pathology.

**Table 2:**
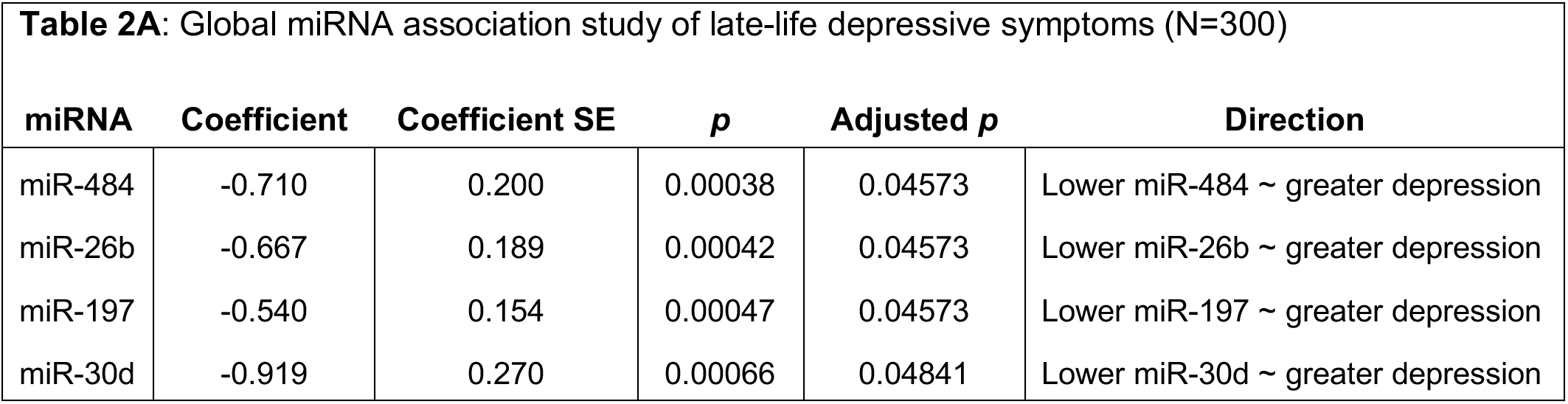
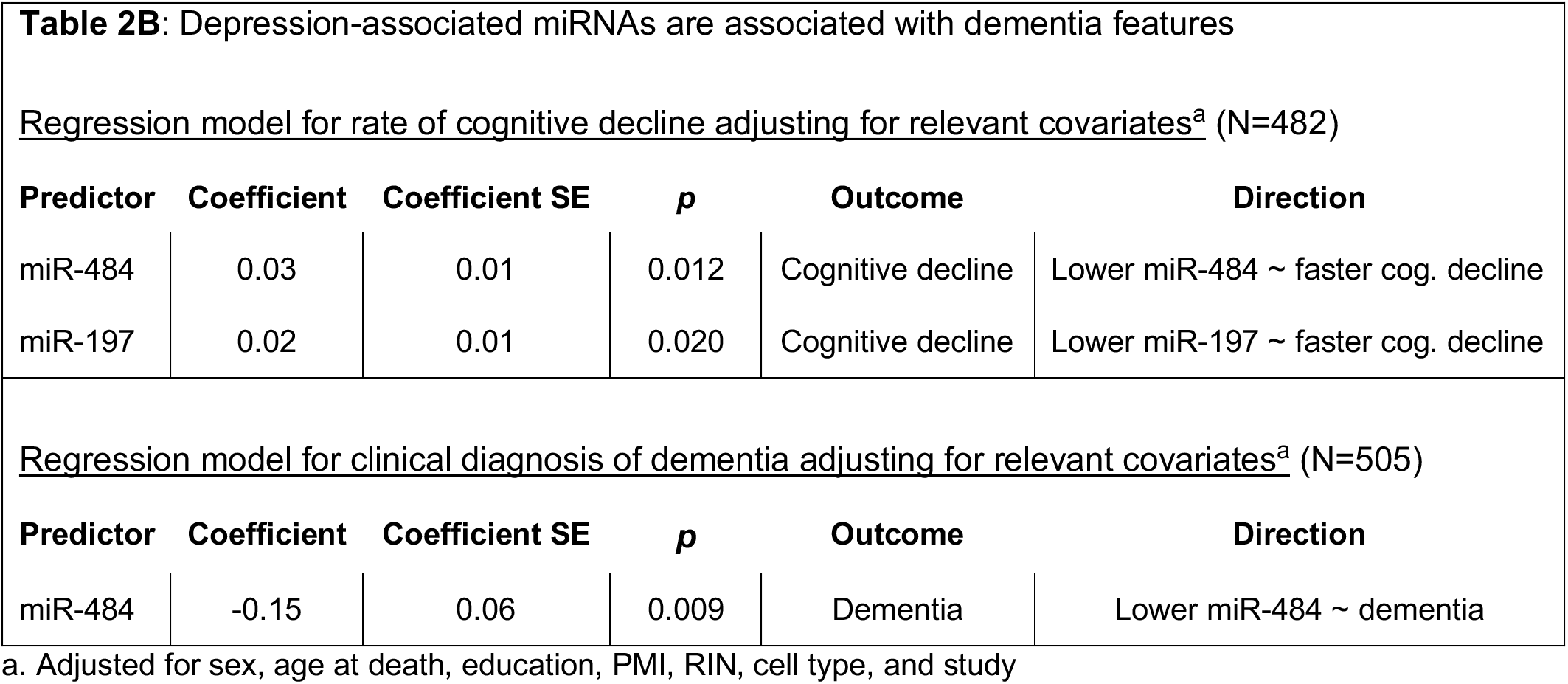
**A)** miRNAs significantly associated with late-life depression at FDR p <0.05 from a global miRNA association study of longitudinally assessed depression, adjusting for sex, age at visit, global AD pathology, proportions of neurons, oligodendrocytes, astrocytes, and microglia, post-mortem interval, RIN, and study. All miRNAs were down regulated in greater depression. **B)** Depression-associated miRNAs are also associated with dementia features after adjusting for sex, age at death, PMI, RIN, proportions of neurons, oligodendrocytes, astrocytes, and microglia, post-mortem interval, RIN, and study.

**Figure 1:**
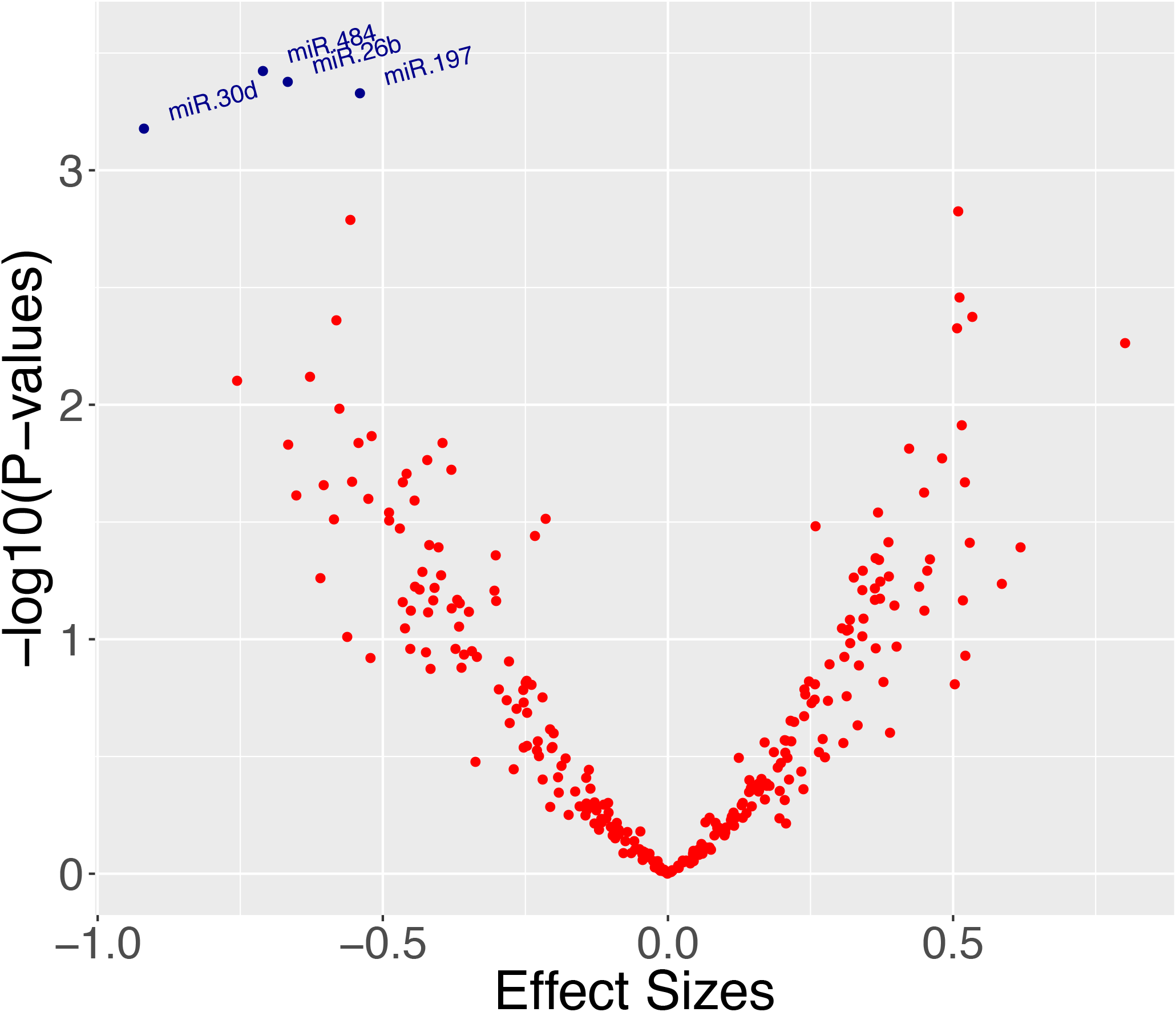
Volcano plot for global miRNA analysis of late-life depression assessed longitudinally. Four miRNAs were significantly associated with late-life depression after adjusting for sex, age at visit, global AD pathology, proportions of neurons, oligodendrocytes, astrocytes, and microglia, PMI, RIN, study, and batch at adjusted p <0.05. These are miR-484, miR-26b, miR-30d, and miR-197.

### Depressive symptoms-associated miRNAs versus dementia features

Given the observation that late-life depression is associated with increased incidence of dementia in several epidemiological studies (1, 2), we examined associations between these four depressive symptoms-associated miRNAs and the features of dementia to test whether these miRNAs might share a common molecular mechanism with dementia. Interestingly, two miRNAs (miR-484 and miR-197) were associated with the rate of cognitive decline over time after adjusting for sex, age at death, education, PMI, RIN, proportions of neurons, oligodendrocytes, astrocytes, microglia, and study (miR-484, β=0.03, p=0.012, Table 2B; miR-197, β=0.02, p=0.020, Table 2B, N=482). Both miR-197 and miR-484 were down-regulated in faster cognitive decline, as well with greater depressive symptoms, consistent with the known detrimental effects of depression on cognitive aging.

Regarding the clinical diagnosis of dementia, miR-484 was significantly associated with clinical diagnosis of mild cognitive impairment or dementia after adjusting for sex, age at death, education, PMI, RIN, cell type proportions, and study (β= −0.15, p=0.009, N=505, Table 2B). Lower miR-484 level was associated with higher odds for MCI or AD, as well as with greater depressive symptoms, which is consistent with the known negative effects of depression on dementia risk.

With regard to dementia pathologies, we examined global AD pathology, gross infarcts, microscopic infarcts, TDP-43, hippocampal sclerosis, cerebral amyloid angiopathy, atherosclerosis, and Lewy bodies in relation to these four miRNAs. However, we did not see a significant association between any of the four depressive symptoms-associated miRNAs and any of the dementia pathologies. These findings are consistent with the observation that depressive symptoms are not directly associated with dementia pathologies (3–6).

Since cognitive decline can be due to neurodegenerative pathologies or independent of them, we examined the associations between miR-484 and miR-197 and cognitive decline while adjusting for all the measured neurodegenerative pathologies (i.e., global AD pathology, gross infarcts, microscopic infarcts, TDP-43, hippocampal sclerosis, cerebral amyloid angiopathy, atherosclerosis, and Lewy bodies) and sex, age at death, PMI, RIN, study, and cell type proportions. Using this model, we found that only miR-197 was significantly associated with cognitive decline (miR-197, β=0.023, p=0.016, N=390). This suggests that miR-197 contributes to cognitive decline via mechanisms independent of the known dementia pathologies and may be a target for further studies of cognitive resilience.

### Enrichment of targets of miR-484, miR-197 in protein co-expression modules

Since a miRNA represses expression of its target genes by either destabilizing their transcripts or repressing the translation of the transcripts to proteins, the effects of miRNAs are likely best observed at the protein level. Hence, we asked whether the targets of miR-484 or miR-197 would be enriched in any of the 16 published protein co-expression modules derived from the proteomes of the Baltimore Longitudinal Study of Aging cohort (26). We found that targets of miR-197 were not enriched in any of these modules. Interesting, the targets of miR-484 were enriched in the M1 turquoise module, which is involved in synaptic transmission and regulation of long-term neuronal synaptic plasticity (enrichment p=0.002; BH adjusted p = 0.034; Figure 2) (26) (26). There were 46 genes that were common between the predicted targets of miR-484 and module M1 turquoise (Supplementary Table 2).

**Figure 2:**
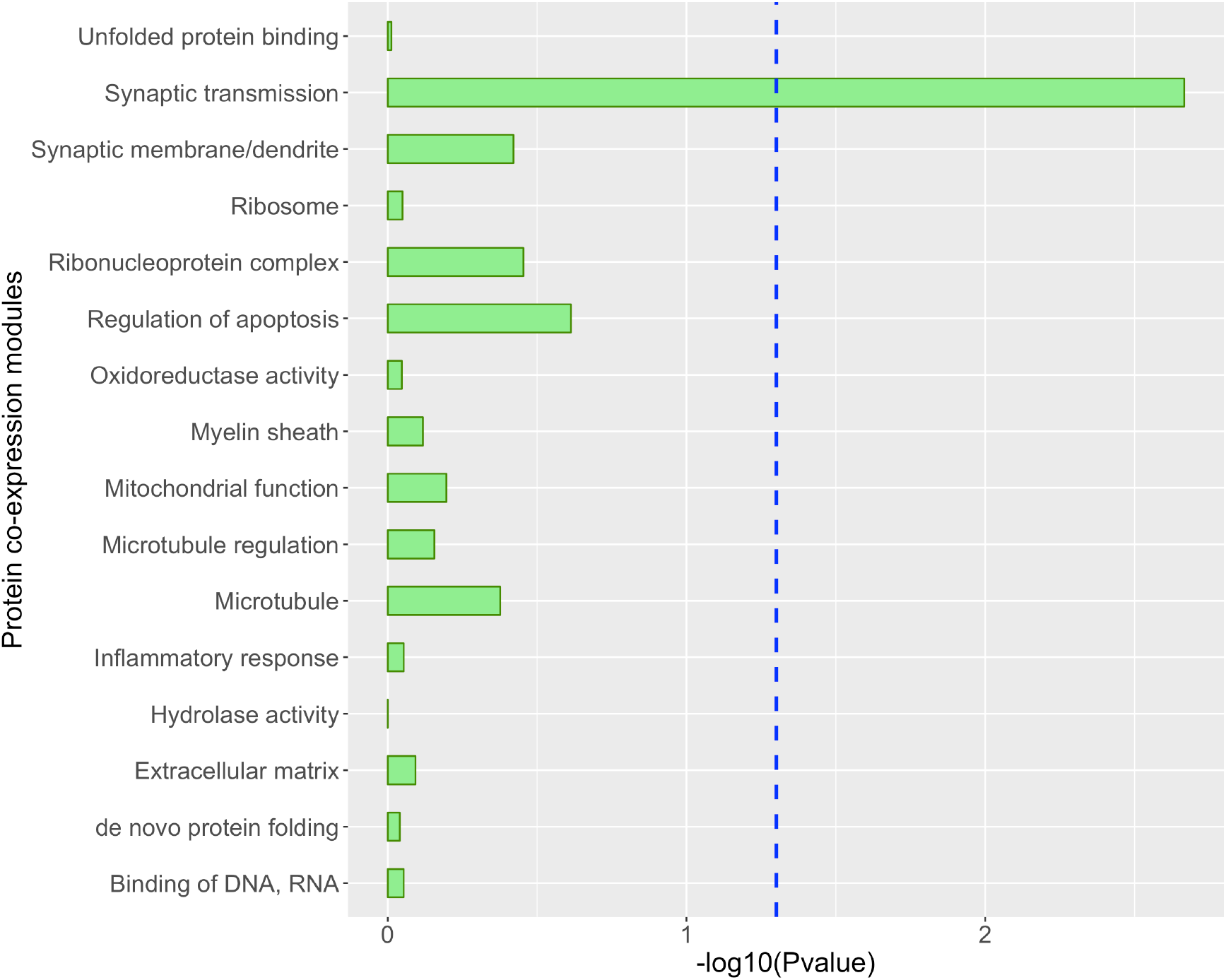
Enrichment of the predicted targets of miR-484 in BLSA protein co-expression modules. The y-axis lists the 16 brain protein co-expression modules and their enriched biological activities from Seyfrieds et al 2017. The blue line perpendicular to the x-axis represents Benjamini-Hochberg adjusted p <0.05.

## Discussion

This is the first global brain miRNA association study of late-life depressive symptoms assessed longitudinally over a mean of seven years in two community cohorts to the best of our knowledge. We found that four miRNAs were significantly down-regulated in greater depressive symptoms (miR-484, miR-197, miR-26b, and miR-30d). Furthermore, among these depressive symptoms-associated miRNAs, miR-484 and miR-197 were significantly down-regulated in faster decline of cognitive performance over time, and miR-197 was associated with faster decline of cognitive performance independent of the eight measured traditional neuropathologies. In addition, lower expression of miR-484 was also significantly associated with higher probability for having a clinical diagnosis of mild cognitive impairment or Alzheimer’s dementia. Taken together, these findings suggest a molecular link between late-life depressive symptoms and elevated risk for Alzheimer’s dementia through miR-484 and miR-197. Interestingly, the predicted targets of miR-484 were enriched in the protein co-expression module involving synaptic transmission and regulation of long-term neuronal synaptic plasticity. We hypothesize that miR-484 acts through its effects on synaptic transmission and neuronal synaptic plasticity in elevating risk for dementia, which should be tested in model systems.

Down-regulation of prefrontal cortex miR-484 and miR-197 in greater depressive symptoms in our data is consistent with results from two prior studies of bipolar disorder, a syndrome predominated by severe depressive symptoms, that observed down-regulation of prefrontal cortex miR-484 and miR-197 in bipolar disorder (28). Bipolar disorder and depression share at least two features. First, individuals with bipolar disorder are depressed the majority (70% to 81%) of their ill-time (29). Second, there is a significant shared heritability between depressive symptoms and bipolar disorder (r = 0.35) (30). Furthermore, down-regulation of prefrontal cortex miR-484 in Alzheimer’s dementia versus control in our study is consistent with the down-regulation of temporal cortex miR-484 in Alzheimer’s dementia relative to controls in a prior study by Pichler and colleagues (31). It is also worth noting that our finding of down-regulation of all the significant miRNAs in late-life depressive symptoms in the prefrontal cortex is consistent with a prior observation of an overall down-regulation of miRNAs in the prefrontal cortex of depressed suicide subjects (8, 11).

miR-484 is known to regulate multiple biological processes including neurogenesis and mitochondrial fission (32–34). miR-484 promotes neurogenesis by inhibiting protocadherin-19 (32). Expression level of miR-484 was found in our study to be reduced in persons having more depressive symptoms, or faster decline of cognitive performance, or a clinical diagnosis of dementia; hence, by inference, these persons likely have reduced neurogenesis. In addition, miR-484 regulates mitochondrial fission, which is a quality control mechanism to protect against oxidative damage in neurons (35). Moreover, altered balance of mitochondrial fission and fusion has been shown to be a an important mechanism leading to mitochondrial and neuronal dysfunction in Alzheimer’s dementia brain (36, 37).

Our study is potentially limited due to the lack of a suitable replication cohort with global miRNA in the dlPFC and measures of late-life depressive symptoms. Hence, associations between these four miRNAs and late-life depressive symptoms need to be replicated in an independent sample. However, this concern is alleviated by the finding of down-regulation of prefrontal cortex miR-484 and miR-197 in bipolar disorder, which shares important features with depression. Our study has several strengths. First, it is the first brain miRNA association study of late-life depressive symptoms to the best of our knowledge. Second, in our study, late-life depressive symptoms was assessed annually and longitudinally over a mean of seven follow-up years. Third, this is the first study to observe a link between late-life depressive symptoms and dementia at the molecular level, via miRNAs. Fourth, we rigorously controlled for potential confounding factors in our analyses including post-mortem brain interval and tissue heterogeneity.

In conclusion, we here show that late-life depressive symptoms is linked to the down-regulation of prefrontal cortex miR-484, miR-197, miR-26b, and miR-30d. Furthermore, the association between late-life depression and elevated risk for dementia observed in 23 prospective epidemiological studies (1, 2, 38) may be partially explained by miR-484 through its effects on synaptic transmission, synaptic plasticity, neurogenesis, and mitochondrial fission. Further mechanistic studies are needed to confirm these observations.

